# Development and Characterization of Chicken Lung Organoids for Future *In Vitro* Modeling of Avian Influenza Virus-Host Cell Interaction

**DOI:** 10.1101/2025.03.12.640685

**Authors:** Hannah Nicholson, Christopher Zdyrski, Megan P. Corbett, Anice C. Lowen, Michael Catucci, Bryan J. Melvin, Lisa J. Stabler, Eugene Douglass, Jonathan P. Mochel, Karin Allenspach, Silvia Carnaccini

**Affiliations:** Precision One Health Initiative, College of Veterinary Medicine, University of Georgia, Athens, GA, USA; Department of Microbiology and Immunology, Emory University School of Medicine, Atlanta, GA, USA; Emory Center of Excellence for Influenza Research and Response (Emory-CEIRR), Atlanta, GA, USA; Population Health, College of Veterinary Medicine, University of Georgia, Athens, GA, USA; Pharmaceutical and Biomedical Sciences, Institute of Bioinformatics, University of Georgia, Athens, GA, USA; Veterinary Diagnostic and Production Animal Medicine, College of Veterinary Medicine, Iowa State University, Ames, IA, USA

**Keywords:** avian, organoids, lung, chicken, single-nuclei RNA seq, stem cells

## Abstract

High pathogenicity avian influenza viruses pose a growing threat to poultry, livestock, wildlife, and humans as they undergo accelerated expansion of host and geographical ranges. Since 2020, these viruses have driven a panzootic characterized by extensive viral diversification and spillover into species previously considered to be resistant to the disease. There is currently a lack of physiologically relevant *in vitro* models that can be used to screen the rapidly changing viral landscape. To address this need, we describe the first chicken lung organoids derived from adult stem cells of specific pathogen free White Leghorns. We analyze their gene expression with bulk RNA sequencing, confirm their cellular heterogeneity via single-nuclei RNA sequencing, and provide basic morphological characterization using hematoxylin and eosin staining, immunohistochemistry, immunofluorescence, and transmission electron microscopy. The results indicate that the organoids contained several cell types, including non-ciliated columnar, cuboidal, squamous, and mucin-producing cells, representative of different regions of the avian respiratory system. Furthermore, expression of genes relevant to influenza A virus infection and replication appeared to be conserved across different sample types. These organoids have the potential to effectively and efficiently model viral infection of the chicken lung, enabling the investigation of viral pathogenesis and evolutionary potential, virus-host interactions, and discovery of targets for antiviral treatments.

## Introduction

High Pathogenicity Avian Influenza Viruses (HPAIVs) have become a serious worldwide threat due to their pandemic potential, devastating effects on poultry and livestock industries, and mortality in wildlife populations. From 2003 to January 2025, 964 cases of human infection with HPAIV (H5N1) have been reported from 24 different countries, with fatal cases (466 total) typically being associated with exposure to infected poultry (WHO, 2025). In the United States, the first recorded human infection of avian influenza occurred in 2022, and the first human death was reported in early January 2025 (CDC, 2025; Garg et al., 2024; LDH, 2025). Since 2020, H5 subtype HPAIV strains of clade 2.3.4.4b have been responsible for mortality in an increasing number of avian and mammalian species, including waterfowl, raptors, cats, ruminants, and sea lions (Adlhoch et al., 2023; Burrough et al., 2024; Chothe et al., 2025; Naraharisetti et al., 2025; Nemeth et al., 2023; Neumann & Kawaoka, 2024; Peacock et al., 2024; Ulloa et al., 2023).

Increased spillover events in mammals, along with recently recognized mammal-to-human transmission, suggest that 2.3.4.4b clade H5N1 viruses have increased potential to cross species barriers relative to ancestral clades, raising concern about their pandemic potential (de Vries & de Haan, 2023).

AIV infections in chickens can occur via the respiratory route, and while in Low Pathogenicity AIV replication is restricted to epithelial cells of the upper respiratory tract, HPAIVs infect both epithelial and non-epithelial cells throughout both the lung and systemically (Rebel et al., 2011). The avian lungs are composed of bronchi, parabronchi, and air sacs, with epithelial cell morphology and function varying between these regions. In the bronchi, the mucosa is lined primarily by pseudostratified ciliated columnar cells, as well as non-ciliated columnar, cuboidal, and goblet cells. The parabronchi are smaller tubular airways arranged in a honeycomb-like network within the avian lung. They are lined by non-ciliated cuboidal to squamous cells and are surrounded by blood capillary-rich air capillaries, where oxygen and carbon dioxide exchange occurs (Lopez- & Cuesta-and Burrell, 2000; Maina, 2015). While 2D cell cultures have been used to study HPAIVs, they lack cellular heterogeneity and the structural complexity of respiratory tissue, both of which are crucial for understanding viral pathogenesis (Bertrams et al., 2022; König et al., 2010; A. Zhou et al., 2021). Therefore, novel *in vitro* methods to study these virus-host interactions may provide insights that are difficult to replicate in current models.

One model that shows promise for studying these complex interactions is adult stem cell-derived (ASC) organoids. ASC-derived organoids are composed of various differentiated epithelial cell types and are established from stem cells that are specific to the parent tissue (Mochel et al., 2017). Organoids are more complex than conventional primary cell cultures due to their composition of multiple epithelial cell types, capacity to self-organize into structures that resemble the tissue of origin, and ability to expand without passage limits (Fatehullah et al., 2016; Lehmann et al., 2019). 3D organoids have been derived from a plethora of species including humans, rodents, small animals, farm animals, and reptiles (Beaumont et al., 2021; Chandra et al., 2019; V. Gabriel et al., 2022a, 2024; Post et al., 2020; Zdyrski et al., 2024).

Additionally, they have been derived from nearly all epithelial tissues (Corrò et al., 2020). While ASC-derived organoids are cultured from adult stem cells residing in the tissue of origin and differentiate into cells specific to that epithelial tissue, organoids derived from induced pluripotent stem cells (iPSCs) arise from pluripotent stem cells that have been reprogrammed from various somatic cells. They are then re-differentiated using growth factors to develop into specific cell types of a desired tissue (Fatehullah et al., 2016; Kim et al., 2020; McCauley & Wells, 2017; Z. Zhao et al., 2022). Interestingly, iPSC-derived lung organoids tend to only recapitulate certain portions of the respiratory tract in a functionally fetal state, such as the respiratory epithelium or parenchyma, while ASC-derived organoids can contain representative cells of the mature proximal and distal airways (Han et al., 2022; Miller et al., 2019; Tindle et al., 2021). Moreover, ASC-derived organoids are primarily epithelial, while iPSC-derived organoids may also include additional cell types, such as mesenchymal and endothelial cells (Kim et al., 2020; Miller et al., 2019).

Despite avian influenza being a major panzootic concern, the poultry industry is also affected by other respiratory illnesses caused by viruses and bacteria, which can result in significant economic losses for producers (Mehrabadi et al., 2022; USDA, 2025). Some of these illnesses, which can involve the infection of epithelial lung cells, include viral Newcastle and infectious bronchitis virus, in addition to bacteria such as *Escherichia coli* and *Chlamydia psittaci* (Boodhoo et al., 2016; Glisson, 1998; Lagae et al., 2016; W. Liu et al., 2023; Mol et al., 2019; Yehia et al., 2023). Therefore, chicken lung organoids have the potential to better model important poultry diseases *in vitro* compared to conventional 2D cell cultures. To the best of our knowledge, chicken organoids have so far been developed only from intestinal crypts with the aim of studying poultry gut health (D. Zhao et al., 2022).

This manuscript documents the establishment, maintenance, and characterization of the first chicken lung organoids derived from adult stem cells. The establishment of this model may advance our understanding of the tropism and replication dynamics of high pathogenicity avian influenza viruses and other respiratory pathogens that impact the poultry industry. Additionally, the successful establishment of cultures from domestic chickens may aid in producing and expanding organoids from other avian species of significance to One Health.

## Methods

### Tissue collection

Three-week-old, unsexed, specific pathogen free White Leghorn chickens (*Gallus gallus domesticus*) purchased from Boehringer Ingelheim were used to derive lung organoids (Gainesville, Georgia, USA). Following AVMA guidelines, the birds were anesthetized using isoflurane and euthanized via cervical dislocation (IACUC: A2023 07-018-A6). In a class II, type A2 biosafety cabinet, both lungs from each donor were dissected and washed in sterile PBS (Corning; 21-040-CM) three times. A piece of tissue from each lung was submerged in a tube containing Shipping Media (composition listed in **Supplemental Table 1**) on ice. A subset of the tissue was flash-frozen in liquid nitrogen for 30 seconds before being stored in a -80°C freezer for future analysis. Additionally, tissue from each donor was fixed in 10% neutral-buffered formalin for 24 hours before being transferred to 70% ethanol for further histopathological processing.

### Development and maintenance of organoid cultures

Methods for establishing and maintaining the organoids were developed from previous work, with minor modifications (Zdyrski et al., 2024). To isolate adult stem cells for organoid culture, tissue from each chicken lung was minced in a petri dish, 6 mL of DMEM/F12 (Gibco; 12634- 010) was added, and all liquid and tissue fragments were collected and transferred to a 15 mL tube. The sample was centrifuged at 100 x g (4°C) for 5 minutes. The supernatant was removed, 500 µL of Red Blood Cell Lysis Buffer (Roche; 11814389001) was mixed with the pellet, and the sample was placed on a nutator at medium speed for 5 minutes. Then, 6 mL of DMEM/F12 was added to the tube, which was spun as before, the supernatant was removed, and the pellet containing tissue was resuspended in Matrigel matrix (Corning; 356255). A 24- well plate (Corning; 3524) was prewarmed at 37°C for 20 minutes before adding 30 µL of the suspension to each well, and the plate was incubated (PHCBi; MCO-170AICUVL) at 37°C and 5% CO^2^ for 20 minutes before adding media (composition found in **Supplemental Table 2**). Media was removed and replenished every Monday (500 µL), Wednesday (500 µL), and Friday (750 µL). To clean (replace Matrigel or adjust density) or passage (dissociate) the sample, organoids were removed from the Matrigel matrix by first discarding the media, adding 500 µL of Cell Recovery Solution (Corning; 354270) to each well, physically disrupting the Matrigel dome with a pipette tip, and transferring all contents to a 15 mL tube. The sample was then incubated on ice for 10 minutes and centrifuged at 100 x g (4°C) for 5 minutes. The supernatant was then removed, leaving ∼500 µL of Cell Recovery Solution. When cleaning, 6 mL of DMEM/F12 was added, followed by centrifugation at the same setting as described above. However, when passaging, 500 µL of TrypLE™ Express (Gibco; 12604-021) was added, mixed with the cell pellet, and incubated in a heat bath (37°C) for 10 minutes, shaking the tube at the 5-minute mark. After pipette mixing, 6 mL of DMEM/F12 was transferred to the tube before centrifuging. For both cleaned and passaged samples, all supernatants were removed. The cell pellet was resuspended in Matrigel (30 µL per well) and plated in pre-warmed 24-well plates. After incubating for 20 minutes at 37°C and 5% CO^2^, media was added to each well. Organoids were imaged with an ECHO Revolution Microscope, and scale bars were added with ImageJ (version 1.54g).

### Cryopreservation and thawing

To cryopreserve the organoids, the cleaning protocol, as described in the previous section, was followed. Then, the cell pellet was resuspended in 1 mL of Cryostor (Biolife Solutions; 210102), transferred to a cryovial, and placed in a Mr. Frosty container (Nalgene; 5100-0001) at 4°C. After 10 minutes, the container was moved to a -80°C freezer for 24 hours before the vials were stored in vapor phase liquid nitrogen (-196°C) indefinitely. To thaw the organoids from long-term storage, vials were recovered from liquid nitrogen and immediately placed in a 37°C bead bath for ∼3 minutes or until thawed. As soon as the sample was thawed, the contents of the cryovial were transferred to a 15 mL tube containing 6 mL of DMEM/F12. The sample was then spun, resuspended in Matrigel, and plated as described above.

### Histology and immunohistochemistry

Media was removed from organoid samples and replaced with 500 µL of formalin acetic acid for 24 hours (V. Gabriel et al., 2022b). Formalin acetic acid (organoids) and formalin (tissues) were replaced with 70% ethanol after this time. Next, tissues were transferred to cassettes. For the organoid samples, entire intact Matrigel droplets were transferred to cassettes with biopsy sponges, and the samples were paraffin embedded. Slides were stained with hematoxylin and eosin (H&E), Alcian Blue pH 2.5, pan-cytokeratin AE1/AE3 (Cell Marque; 313M-16), and TTF-1 (Cell Marque; 343M-96) at the University of Georgia Histology Laboratory. Tissue H&E slides and all IHC slides were imaged with an Olympus BX41 Microscope with a BioVid camera.

Images were color-corrected, and scale bars were added with ToupView, version x64 4.11.23945.20231121 (LW Scientific). For higher magnification, H&E organoid slides were imaged with an Olympus BX41 Microscope using the Olympus DP71 camera. Images were color-corrected, and scale bars were added with the Olympus cellSens Standard 1.16 (Build 15404) software.

### Immunofluorescence

Indirect immunofluorescence staining was performed with monoclonal mouse anti-acetylated tubulin (Millipore; T7451) on both sample types at a 1:1000 dilution. Samples were then stained with secondary antibody, Goat anti-Mouse IgG (H+L) Highly Cross-Adsorbed Secondary Antibody, Alexa Fluor™ Plus 488 (Thermo Fisher Scientific; A32723), and counterstained with Hoechst 33342 (Thermo Fisher Scientific; 62249) at 1:1000 dilutions. All antibodies were diluted in PBS (Sigma Aldrich; 1003434154).

To deparaffinize the sections, slides were sequentially washed in xylene (Fisher Chemical; X3P- 1GAL) three times for 5 minutes each. The process was then repeated, submerging the slides in 100% ethanol. The slides were washed in PBS on an orbital shaker for 5 minutes before performing heat-induced antigen retrieval. Slides were placed on a plastic staining rack in a beaker, and 100X Tris-EDTA Buffer, pH 9.0 Antigen Retrieval Buffer (Abcam; AB93684) diluted to 1X in distilled water was used to fill the beaker enough to cover the entire tissue or organoid section. The slides were microwaved on low power for a total of 10 minutes, stopping at 5 minutes to ensure the buffer solution did not boil over. The slides were removed from the microwave and left at room temperature in the buffer solution for 10 minutes before being washed again in PBS on an orbital shaker for 5 minutes. After drying each slide, a hydrophobic pen (Vector Laboratories; H-4000) was used to draw a border around each section. Using 0.2% Triton X-100 (Thermo Fisher Scientific; A16046.AP)/PBS, 100 µL was pipetted onto the section for 15 minutes before washing in PBS as before. Slides were blocked in 3 drops BLOXALL Blocking Solution (Vector Laboratories; SP-6000) and placed in a humidity chamber filled with PBS at room temperature for 10 minutes. After removing the blocking buffer, 100 µL of primary antibody was added, a piece of Parafilm (Amcor; PM-996) was lightly placed on top of each section, and the slides were left overnight in a humidity chamber at 4°C. The next day, the primary antibody was removed and washed in PBS as previously described. Because the fluorescent antibodies are light-sensitive, the remaining steps were done in the dark. Next, 100 µL of diluted secondary and Hoechst were added, and the slides were incubated in a dark humidity chamber for 60 minutes. After washing as before, one drop of Fluoroshield Mounting Medium (Sigma Aldrich; F6182-20ML) was added to each section, and cover slips (Fisher Scientific; 12541018) were placed on top of each slide before immediately imaging. Immunofluorescent slides were imaged with an ECHO Revolution Microscope. Scale bars were added with ImageJ (version 1.54g).

### RNA extractions

After at least 4 passages, organoids were preserved for bulk RNA-sequencing by following the cleaning protocol described in the development and maintenance of organoid cultures section.

After removing 6 mL of DMEM/F12 supernatant, the cell pellet was resuspended in 100 µL PBS, moved to a cryovial containing 900 µL RNAlater (Invitrogen; AM7021), and stored at -80°C. For RNA extractions, the organoid samples were thawed and transferred to 15 mL tubes, washed with 2 mL of PBS, and centrifuged at 1,200 x g (4°C) for 5 minutes. The supernatant was removed, the cell pellet was resuspended in 1 mL of Trizol (Invitrogen; 15596026), and each sample was briefly vortexed. Flash frozen tissues were transferred from cryovials to microcentrifuge tubes, 800 µL of Trizol were added to each tube, and tissues were pestled until homogenized. Both tissue and organoid samples were left at room temperature for five minutes before centrifuging at 12,000 x g (4°C) for 10 minutes. The supernatant was transferred to a microcentrifuge tube, and 160 µL or 200 µL of chloroform (Alfa Aesar; J67241) were added for tissues and organoids, respectively. The samples were vigorously mixed by shaking for 20 seconds. After sitting at room temperature for 2-3 minutes, samples were spun at 10,000 x g for 18 minutes (4°C). The aqueous top layer was collected and moved to a sterile RNase-free tube before adding an equal amount of 100% RNA-free ethanol. Up to 700 µL were loaded in a Qiagen RNeasy column and collection tube (RNeasy Mini kit; Qiagen; 74104). Samples were centrifuged at 8,000 x g for 30 seconds, the waste in the collection tube was discarded, and DNase treatment was performed according to Qiagen’s protocol (Qiagen; 79254). In a new collection tube, 500 µL buffer RPE was added to the column, and the samples were spun at the same settings as before. After discarding the flow-through, this step was repeated, but samples were now spun for 2 minutes at 8,000 x g. The flow-through was discarded again, and the samples were centrifuged at the same settings for 1 minute. 50 µL of RNase-free water (Sigma; W4502-50ML) was added to the column and incubated for 2 minutes. Samples were centrifuged twice at 8,000 x g for 1 minute. RNA concentration was determined with a Nanodrop ND-1000 Spectrophotometer (Thermo Fisher Scientific), and samples were kept at -80°C until shipped.

### RNA sequencing

All samples were sent to Azenta Life Sciences (South Plainfield, NJ, USA) for preparation of the RNA library and bulk RNA sequencing. A Quibit 2.0 Fluorometer (Life Technologies, Carlsbad, CA, USA) was first used to quantify RNA, followed by assessment of the RNA integrity with an Agilent TapeStation 4200 instrument (Agilent Technologies, Palo Alto, CA, USA). A NEBNext Ultra II Directional RNA Library Prep Kit for Illumina (NEB, Ipswich, MA, USA) was used to prepare the RNA sequencing libraries, using the manufacturer’s instructions. Then, the mRNA was enriched with Oligo(dT) beads and fragmented for 15 minutes (94°C). Both cDNA strands were synthesized, and at the 3’ ends, fragments were end repaired and adenylated. The cDNA fragments were ligated with universal adapters, and limited-cycle PCR was used for index addition and library enrichment. An Aligent TapeStation (Agilent Technologies, Palo Alto, CA, USA) was utilized to confirm the sequencing libraries, and both a Qubit 2.0 Fluorometer (Invitrogen, Carlsbad, CA) and quantitative PCR (KAPA Biosystems, Wilmington, MA, USA) were used for quantification. Sequencing libraries were multiplexed and clustered onto one flow cell. Using the manufacturer’s protocol, an Illumina instrument was then used to sequence the samples with 2x150bp Paired End (PE) configuration. Control Software (HCS) was used for image analysis and base calling. Illumina HiSeq created raw sequence data (.bcl files), which were converted to fastq files and de-multiplexed with Illumina’s bcl2fastq 2.17 software. For index sequence identification, one mismatch was allowed.

### Organoid harvesting for single-nuclei RNA sequencing

Organoids were harvested for single-nuclei RNA sequencing via collection from the Matrigel matrix using the cleaning protocol previously described. Organoids were resuspended in 500 µL DMEM/F12 and transferred to a cryovial. After centrifuging at 100 x g (4°C) for 5 minutes, the supernatant was removed, and the sample was flash-frozen in liquid nitrogen for 30 seconds.

Samples were stored in a -80°C freezer and sent to Genewiz from Azenta Life Sciences (South Plainfield, NJ, USA) on dry ice for analysis.

### Single-nuclei RNA sequencing

Upon arrival, tissue and organoid samples were stored in liquid nitrogen for further processing. Using the manufacturer’s instructions, nuclei were extracted from both samples using Miltenyi Nuclei Extraction Buffer (Miltenyi Biotec, Auburn, CA, USA), in addition to gentle MACS Dissociation and C tubes. Once isolated, AO/PI dye was used on the Nexcelom Cellaca MX to count the nuclei, and a Chromium Single Cell 3’ kit (10X Genomics, CA, USA) was used to create single nuclei RNA libraries. Following the manufacturer’s guidelines, the nuclei were loaded and processed on the Chromium X, targeting approximately 6,000 gel bead-in-emulsions (GEMs) per sample. An Agilent TapeStation (Agilent Technologies, Palo Alto, CA, USA) evaluated quality of the sequencing libraries, and a Qubit 2.0 Fluorometer (Invitrogen, Carlsbad, CA, USA) quantified them. qPCR (Applied Biosystems, Carlsbad, CA, USA) was also used to quantify libraries before loading the samples on an Illumina HiSeq 4000 (or equivalent) instrument. Then, samples were sequenced using the 10X Genomics recommended guidelines. The raw sequence data (.bcl files) were converted to fastq files before being de-multiplexed with the Genomics’ cellranger mkfastq command.

### Bulk and single-nuclei RNA sequencing analysis

Read quality was assessed using FastQC, and high-quality reads were mapped to the galgal5.0.91 transcriptome for single-nuclei RNA sequencing and the GRCg7b genome (Ensembl, 2024). Kallisto or CellRanger was used to conduct read alignment and gene expression counts for bulk and single-nuclei RNA sequencing, respectively (Bray et al., 2016; Zheng et al., 2017). Single-nuclei RNA sequencing data was normalized and log transformed using R package Seurat (Butler et al., 2018; Hao et al., 2021, 2024; Satija et al., 2015; Stuart et al., 2019). R and corresponding R scripts were then used for data preparation procedures, analysis procedures, and visualization (R Core Team, 2021). Data was stored on a GitHub repository, and Zenodo was used to create Digital Object Identifiers (DOIs) for publication citations. Gene Set Enrichment Analysis with the fgsea package was used for pathway analysis, and gene signatures were acquired from the Molecular Signatures Database (MSigDB) (Korotkevich et al., 2021; Liberzon et al., 2011).

### Transmission electron microscopy

Organoids were cleaned as previously described above. After removing supernatant, the cell pellet was resuspended in 1 mL of modified Trump’s electron microscopy fixative (1% paraformaldehyde, 3% glutaraldehyde, .1 M cacodylate), transferred to a cryovial, and stored at 4°C. Samples were then washed in 0.1 M Cacodylate-HCl buffer (pH 7.2) and centrifuged.

Then, the organoids were agar-enrobed in 4% Noble Agar (aqueous), set, and then placed 1% osmium tetroxide in deionized water buffer for 1 hour. Samples were washed in PBS before being placed in 1% osmium tetroxide buffer for one hour. After washing in deionized water, organoids were incubated in 0.5% aqueous uranyl acetate en bloc for one hour in a dark room. The organoids were washed in deionized water again and then dehydrated sequentially using an ethanol series (30%, 50%, 75%, 95%, 100%). Samples were then cleared in two changes of acetone, followed by two changes of propylene oxide before being infiltrated with 2:1, 1:1, and 1:2 mixtures of propylene oxide and Mollenhauer’s Epon-Araldite plastic mixture for two hours, respectively (Mollenhauer, 1963). Samples were changed with 100% Epon-Araldite plastic twice for two hours each. After embedding the samples in flat embedding molds, they were polymerized in a 70-80°C oven overnight (Bazzola, 1992). A Reichert Ultracut S ultramicrotome (Leica, Inc., Deerfield, IL) was used to cut the polymerized blocks into 1 µm sections. After placing on glass slides, the sections were stained with 1% Toluidine Blue O in 1% sodium borate. After choosing areas of interest, the block was trimmed for thin sectioning. Sections (60-70 nm) were obtained and placed on 200-mesh copper Locator grids. Using 2% aqueous uranyl acetate and Reynolds lead citrate, one grid was stained, leaving the other grids unstained (Reynolds, 1963). A JEM-2100 PLUS Transmission Electron Microscope (JEOL, Inc., Tokyo, Japan) with an accelerating voltage of 120 kV was used to view the sample, and the sample was imaged with an AMT NanoSprint 15L-MarkII High Sensitivity sCMOS TEM Camera with a resolution of 5056 x 2960 pixels.

### Contamination testing

Organoid samples were routinely tested for Mycoplasma species contamination with Invitrogen Mycostrip tests (InvivoGen; rep-mys-50), using the manufacturer’s protocol. Additionally, organoids were sent to IDEXX BioAnalytics (Columbia, MO, USA) to confirm negative detection in each cell culture via a Mycoplasma PCR assay. The samples were also confirmed to be of chicken (*Gallus gallus*) origin via IDEXX’s CO1 DNA Barcoding assay prior to cryopreservation. Before IDEXX’s analysis, organoids were cleaned as previously described with minor modifications. The cell pellet was washed in 6 mL PBS, instead of DMEM/F12, twice. After removing the supernatant, the pellet was resuspended in 700 µL PBS and transferred to a cryovial. The sample was centrifuged at 100 x g for 5 minutes at 4°C, the supernatant was removed, and the cryovial was placed in a -80°C freezer until shipped.

## Results

### Organoid growth

Detailed sample information can be found in **Supplemental Table 3**. Organoid growth was observed as early as 24 hours after isolation. Prior to the first passage, cilia were observed on a subset of the organoids, causing them to rotate in the Matrigel. After the first passage, cilia were not visible with brightfield imaging in any sample. On average, samples were passaged every 1-2 weeks (**Figure 1a, 1b, and 1c**). A total of 12 passages were attempted using Chicken Lung 1, totaling 128 days in culture, with no indications of a slowed growth rate (**Supplemental Table 3**). After expansion, all samples were cryopreserved several times, and each organoid line had the capacity to be expanded for multiple passages after thawing (**Supplemental Table 3**). Prior to cryopreservation, all organoid lines were confirmed negative for Mycoplasma spp. using the IDEXX Bioanalytics PCR service. Samples were also confirmed to be of chicken origin via the IDEXX CO1 DNA Barcoding service.

**Fig. 1:**
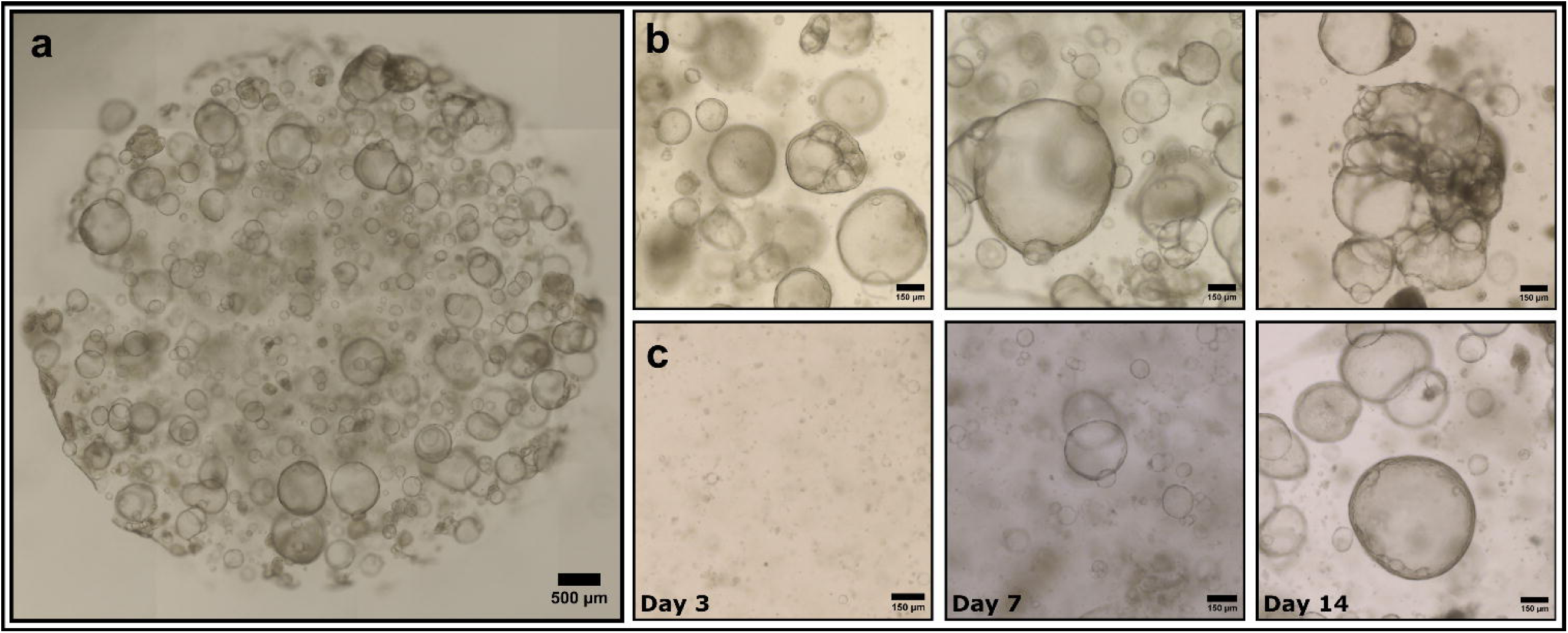
Morphological characterization of chicken lung organoids. (**a**) Whole well scan of the Matrigel droplet containing organoids. (**b**) Representative light microscopy images of three chicken organoid lines. (**c**) Representative microscopy images of an organoid line and regrowth after passage until the next passage. Scale bars are in μm.

### Tissue morphology

Histology confirmed that all lung tissue sections from each donor chicken contained representative structures and cell types expected in the avian lung, from the secondary bronchi to air capillaries. Further, tissues were morphologically intact and did not present any evidence of ongoing clinical disease or microscopic infection. In the tissue sections, no primary bronchi were present. The secondary bronchi, which unlike primary bronchi lack cartilage, were lined primarily by ciliated pseudostratified columnar to cuboidal cells (**Figure 2a**). The secondary bronchi transitioned into the prevalent parabronchi, where air and blood capillaries, lined with simple squamous epithelium and endothelium, respectively, made up the parabronchial wall surrounding the lumen (**Figure 2a and 2c**). Atria, which form the space that led from the parabronchial lumen to the air capillaries, were also found within the walls of the parabronchi. Interatrial septa contained smooth muscle lined with partially ciliated cuboidal cells that transitioned to simple squamous cells (**Figure 2a and 2b**).

**Fig. 2:**
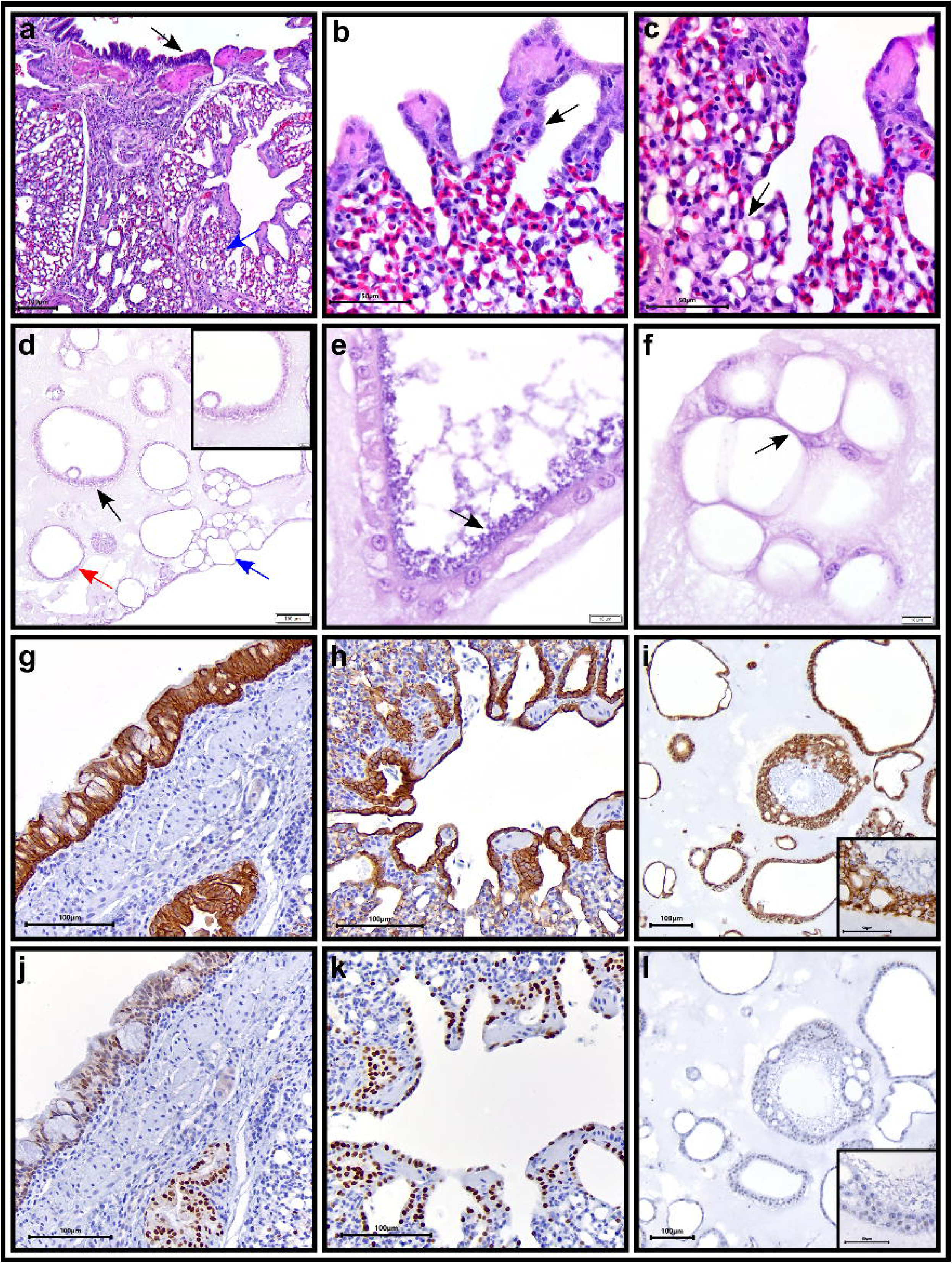
H&E histological and immunohistochemical characterization of chicken lung tissue and organoids. (**a**) Representative image of chicken lung tissue containing pseudostratified columnar respiratory epithelium (black arrow) and parabronchi with air and blood capillaries (blue arrow) making up the parabronchial walls surrounding a lumen. (**b**) Partially ciliated cuboidal epithelium (black arrow) lining the parabronchi and interatrial septa in chicken lung tissue. (**c**) Squamous epithelium lining air capillaries (black arrow) in chicken lung tissue. (**d**) Representative image of organoid culture with pseudostratified (black arrow) to simple low columnar, cuboidal (red arrow), and squamous cells (blue arrow). (**e**) Representative image of mucoid material production (black arrow) in a subset of organoids. (**f**) Squamous epithelial cells in organoids forming air capillary bridges (black arrow). Immunohistochemical staining of (**g**,**h**) chicken lung tissues and (**i**) organoids for pan-cytokeratin. Immunohistochemical staining of (**j**,**k**) chicken lung tissues and (**l**) organoids for thyroid transcription factor-1 (TTF-1). Scale bars are in μm.

Immunohistochemical staining with pan-cytokeratin AE1/AE3 confirmed the morphology and distribution of epithelial cells throughout the lung (**Figure 2g and 2h and Supplemental Figure 1a**). Strong cytoplasmic immunostaining was noted in the ciliated respiratory epithelium of the secondary bronchi (**Figure 2g**). Additionally, cuboidal cells lining the parabronchi and pneumocytes in the air capillaries were immunopositive (**Figure 2h**). Thyroid transcription factor-1 (TTF-1) staining marked bronchial epithelial cells (**Figure 2j and 2k and Supplemental Figure 1b**). While positive nuclear immunostaining was found in epithelial cells throughout the bronchi, parabronchi, and air capillaries, most expression was localized in the nuclei of the epithelial cells of the parabronchi lining the lumen, including type II pneumocytes and secretory club cells (**Figure 2k**). In the respiratory epithelium of the secondary bronchi, positive staining was present, but less prevalent (**Figure 2j**). Immunofluorescent staining identified protein expression of acetylated alpha-tubulin (TUBA4A), a marker for mature cilia, localized on the apical surface of columnar to cuboidal respiratory epithelial cells in the secondary bronchi **(Supplemental Figure 1e and 1f).**

Alcian Blue pH 2.5 was used to stain mucus produced by goblet cells in the respiratory epithelium, in addition to mucoid material in the parenchyma (**Figure 3a and 3b**). TEM revealed lamellar bodies located in the cytoplasm of type II pneumocytes (**Figure 3d and 3f**). Type I pneumocytes lined the air capillaries, and nucleated erythrocytes were identified within the blood capillaries and in the septa of the parabronchi (**Figure 3f and 3g**). In the parabronchi, microvilli were also prevalent on the surface of a subset of cells (**Figure 3d and 3f**).

**Fig. 3:**
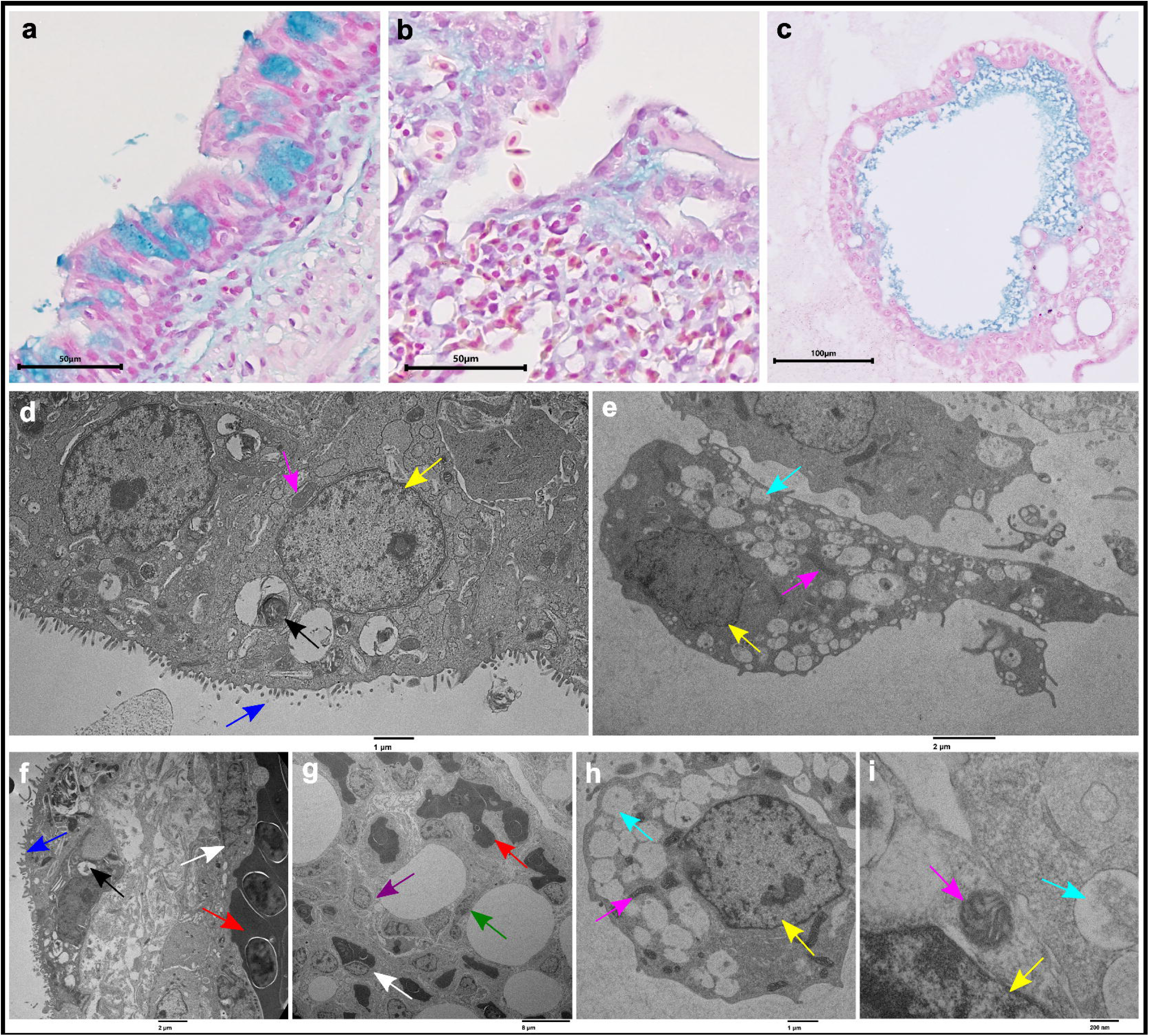
Special staining and TEM characterization of chicken lung tissues and organoids. Representative images of Alcian Blue pH 2.5 staining on (**a,b**) lung tissue and (**c**) organoids. (**d**) Representative TEM micrograph of chicken lung tissue, identifying the nucleus of a cell (yellow arrow), microvilli (blue arrow), mitochondria (pink arrow), and vacuoles containing lamellar bodies (surfactant) (black arrow). (**e**) Representative TEM micrograph of chicken lung organoid with the nucleus (yellow arrow), mitochondria (pink arrow), and vacuoles (teal arrow) highlighted. (**f**) TEM micrograph of chicken lung tissue, including microvilli (blue arrow), surfactant (black arrow), endothelial cells (white arrow), and erythrocytes (red arrow). (**g**) TEM micrograph of the air capillaries in the lung tissue, including type I pneumocytes (green arrow), type II pneumocytes (purple arrow), erythrocytes (red arrow), and endothelial cells (white arrow). (**h,i**) TEM micrographs of organoid cells, identifying nuclei (yellow arrow), mitochondria (pink arrow), and vacuoles (teal arrow). Scale bars are in μm and nm.

### Organoid morphology

Using a light microscope, organoids were identified as primarily large and round, with a dark border and hollow lumen (**Figure 1a and 1b**). A smaller subset of organoids seemed to contain multiple hollow and thin-walled, cystic structures (**Figure 1b)**. Organoids in each culture displayed a variety of epithelial cell types observed in H&E staining. These included low columnar, cuboidal, low cuboidal, and squamous cells, likely formed after partial differentiation from adult stem cells and resembling the epithelia of the bronchi, parabronchi, and air capillaries, respectively. Most organoids were composed of a simple epithelium surrounding a large lumen (**Figure 2d**). However, there were subsets containing pseudostratified epithelium, which in some areas appeared to undergo squamous metaplasia (**Figure 2d**). H&E staining revealed basophilic mucoid material within lumens formed by pseudostratified to simple cuboidal and columnar cells (**Figure 2e**). Alcian Blue pH 2.5 staining confirmed the production of acidic mucins, indicative of goblet-like cells (**Figure 3c**).

Interestingly, simple squamous cells appeared to form interconnected bridges (**Figure 2f**), resembling the air capillary bridges found *in vivo*. Strong cytoplasmic pan-cytokeratin immunostaining indicated that most cells in culture were epithelial by nature, apart from some presumed to be fibroblasts (**Figure 2i and Supplemental Figure 1a**). TTF-1 immunohistochemical staining identified the bronchial-like epithelial cells (**Figure 2l and Supplemental Figure 1b**). In the organoids, some cuboidal and squamous cell nuclei were faintly immunopositive compared to the lung tissue.

TEM micrographs revealed many vacuoles and an abundance of mitochondria in the cytoplasm of the organoids, resembling those found in a subset of tissue cells (**Figure 3e, 3h, and 3i**). Of the observed cells, there was no indication of lamellar bodies or surfactant (**Figure 3e, 3h, and 3i**). Additionally, cilia were not observed on any of the cells analyzed, and TUBA4A immunofluorescent staining was negative (**Supplemental Figure 1g and 1h**).

### Bulk RNA sequencing

Tissues and organoids from all samples were sequenced in triplicate. The fidelity of each organoid sample compared to its parent tissue was assessed. A principal component analysis (PCA) plot on internally normalized organoids and tissue samples revealed strong correlation between samples of each donor, indicating that overall gene expression in the parent tissue and organoids was highly conserved (**Figure 4a**). Sample correlation showed that tissue and organoid samples clustered separately, and replicates from each donor clustered together (**Figure 4b**). Overall, there were 11,473 shared genes between tissues and organoids, with 1,203 distinct genes in tissues and 64 distinct genes in organoids (**Figure 4c**). Furthermore, there was a high correlation between organoid and tissue gene expression (**Figure 4d**).

**Fig. 4:**
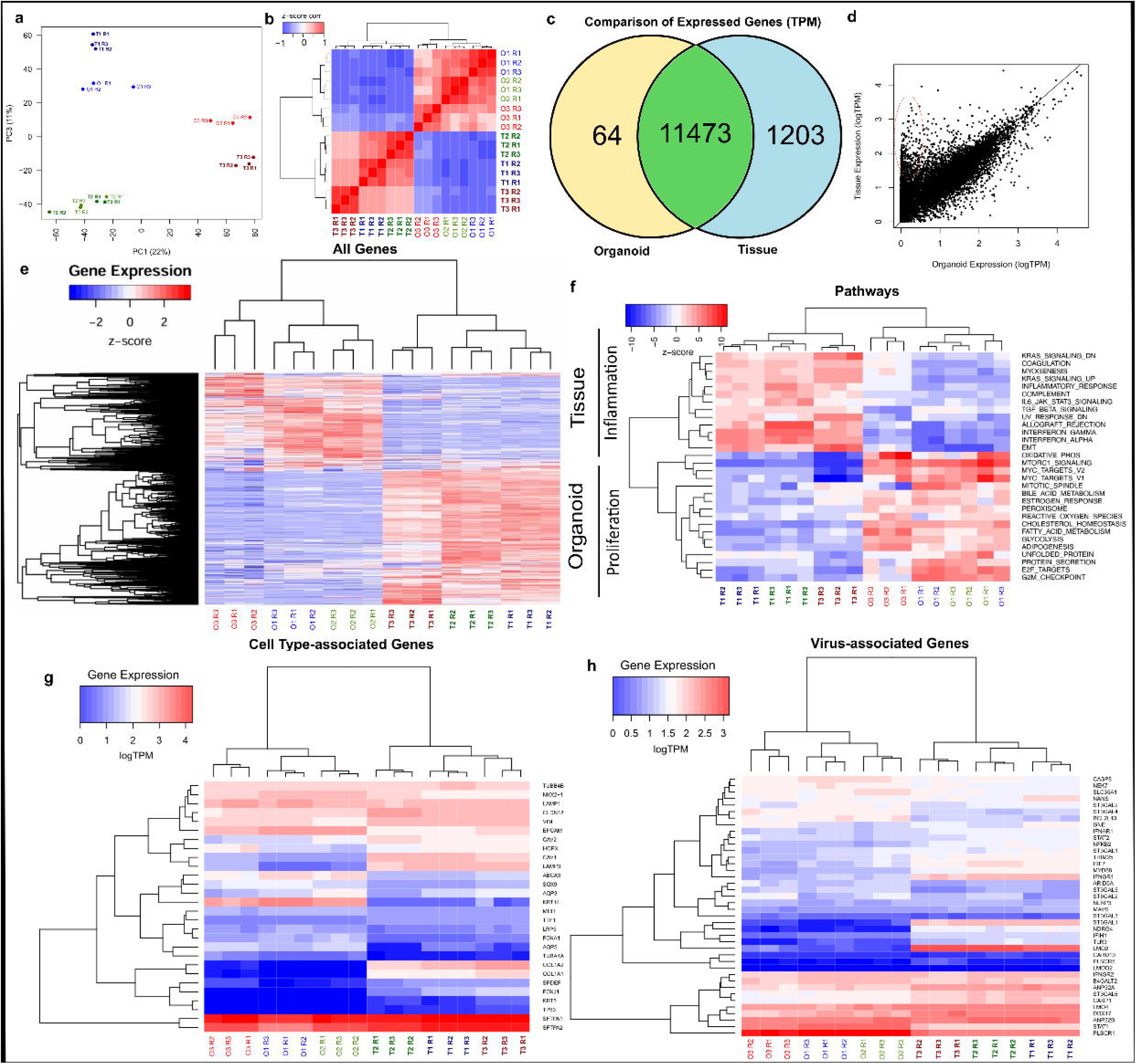
Bulk RNA-Seq analysis of chicken lung organoids and their tissue of origin. (**a**) PCA plot of organoid samples with their parent tissues based on internal normalization of all tissue and organoid samples. Three donors are displayed: Chicken 1 (blue), 2 (green), and 3 (red) with each sample sequenced in triplicates (n = 3 technical replicates). Tissue samples are bolded. (**b**) Sample correlation plot of organoids and tissues. (**c**) Venn diagram comparing genes expressed in both sample types, and those unique in tissues and organoids (**d**) Scatter plot revealing high correlation of organoid and tissue gene expression, in addition to a subset of tissue-specific genes, denoted in red. (**e**) Heatmap of all genes relatively expressed in tissue and organoid samples (upregulation = red, neutral = white, downregulation = blue). (**f**) Heatmap of major pathway analysis, mainly consisting of inflammatory-related and proliferative-associated pathways. (**g**) Heatmap containing genes of interest for epithelial and mesenchymal cell types of the lung and their expression in tissues and organoids. (**h**) Heatmap displaying expression of viral-associated genes and comparison of expression between tissue and organoid samples. Heatmap scales are in z-score (**e,f**) and logTPM (**g**,**h**).

Comparing relative expressions in tissue and organoid samples based on z-score, tissues and organoids did cluster separately overall (**Figure 4e**). Pathway analysis of the tissue vs. organoid samples revealed that in general, the tissues predominantly expressed pathways related to inflammation due to the presence of inflammatory immune cells, while the organoids expressed mostly pathways related to proliferation (**Figure 4f**).

Next, several genes of interest associated with specific cell types were mapped. While organoids clustered separately to tissues, trends in absolute expression were largely conserved across samples, emphasizing the similarities of the organoid cultures and parent tissues (**Figure 4g**). These trends were likely driven by the fact that the organoids are comprised of epithelial cells and therefore exhibit higher expression of genes associated with epithelial cells than mixed tissue samples. Tissues and organoids similarly had high expression of *SFTPA1* and *SFTPA2* which are both associated with type II pneumocytes. *ABCA3*, also associated with type II pneumocytes, was more highly expressed in organoids compared to tissues. Both sample types expressed *AQP5*, a marker for type I pneumocytes, at similar levels.

Epithelial-related genes including *EPCAM* and *KRT14* were upregulated in the organoids, while *KRT5* showed higher expression in the tissue. Genes associated with mesenchymal cells, including *VIM*, *COL1A1*, and *COL1A2*, had increased expression in the tissues as expected because ASC-derived organoids are mainly epithelial. Genes involved in ciliated cell lineage, such as *TUBA4A* and *TUBB4B* were both present in organoids and tissues with similar TPM. Interestingly, *TUBB4B* expression was higher in both tissues and organoids compared to *TUBA4A*. When looking at *TUBA4A* expression between tissues and organoids, the organoids showed greater expression.

Additionally, many viral-associated genes were present in both sample types (**Figure 4h**). *IFIH1*, the precursor for melanoma differentiation associated gene–5 (MDA5), was present in both organoids and tissues. Furthermore, a subset of genes associated with sialic acids, which are bound by hemagglutinin proteins of influenza viruses, were present in the organoids (Van Poucke et al., 2010; C. Zhao & Pu, 2022). Similar transcript numbers were found in organoids and tissues for the genes including: *TLR3*, *TRIM25*, *CASP1*, *NEK7*, and *NLRP3*. These genes are associated with the NLRP3 inflammasome signaling pathway, which is an important part of innate immunity implicated during the host’s initial response to a virus (Kuriakose & Kanneganti, 2017). Furthermore, genes associated with the NF- κB pathway were also present, including *MYD88*, *CARD10*, and *NFKB2* in both tissues and organoids (Chen et al., 2022). *ANP32A* and *ANP32B*, which are host cell genes required for the replication of avian influenza viruses, were also present in both sample types (Long et al., 2019; Staller et al., 2019). Additionally, *PLSCR1*, which is associated with inhibiting viral replication, was more highly expressed in organoids versus tissues (Y. Liu et al., 2022).

### Single-nuclei RNA sequencing comparison between tissue and organoids

Both tissue and organoids (after passage 4) from Chicken #3 were sequenced via single-nuclei RNA seq and mapped to the galgal5.0.91 transcriptome. A total of 3,711 cells were sequenced in the tissue and 18,311 cells in the organoids. Most reads mapped to the chicken genome with 86.4% in tissues and 90.1% in organoids. In the tissue, there were 17,012 total genes detected, with 100,087 mean reads per cell and 644 median genes per cell. In the organoids, there were 18,369 total genes detected, 39,325 mean reads per cell, and 1,834 median genes per cell. In the tissue, a total of 9 distinct cell clusters by a uniform manifold approximation and projection (UMAP) were identified as epithelial, pneumocyte, ciliated, endothelial, smooth muscle, fibroblasts, erythrocytes, immune, and proliferative cells (**Figure 5a**). The cluster of proliferative cells could not be resolved into more specific cell types, and these cells were also present in the epithelial, fibroblast, endothelial, and immune cell clusters (**Figure 5a**). Additionally, while this cluster highly and uniquely expressed *NDC80*, *CEP55*, *CENPF*, *KIF23*, and *TOP2A*, a similar gene expression was shared with endothelial cells (**Figure 5d and 5f**). Interestingly, there were two distinct populations of fibroblasts within the same cluster, and a subpopulation was characterized as perivascular fibroblasts (**Figure 5a**). There were 3 unique clusters found in the organoids: epithelial, cycling epithelial, and fibroblasts (**Figure 5b**). In addition to the same genes highly expressed in the epithelial cell cluster, cycling epithelial cells uniquely had high expression of *CENPP*, *DIAPH3*, *BRCA1*, *CIT*, and *SLX4IP* (**Figure 5d and 5f**). The fibroblast cell cluster uniquely had higher expression of *EPHA3*, *PREX2*, *ITGA8*, *NAV3*, and *EFEMP1* **(Figure 5d and 5f).**

**Fig. 5:**
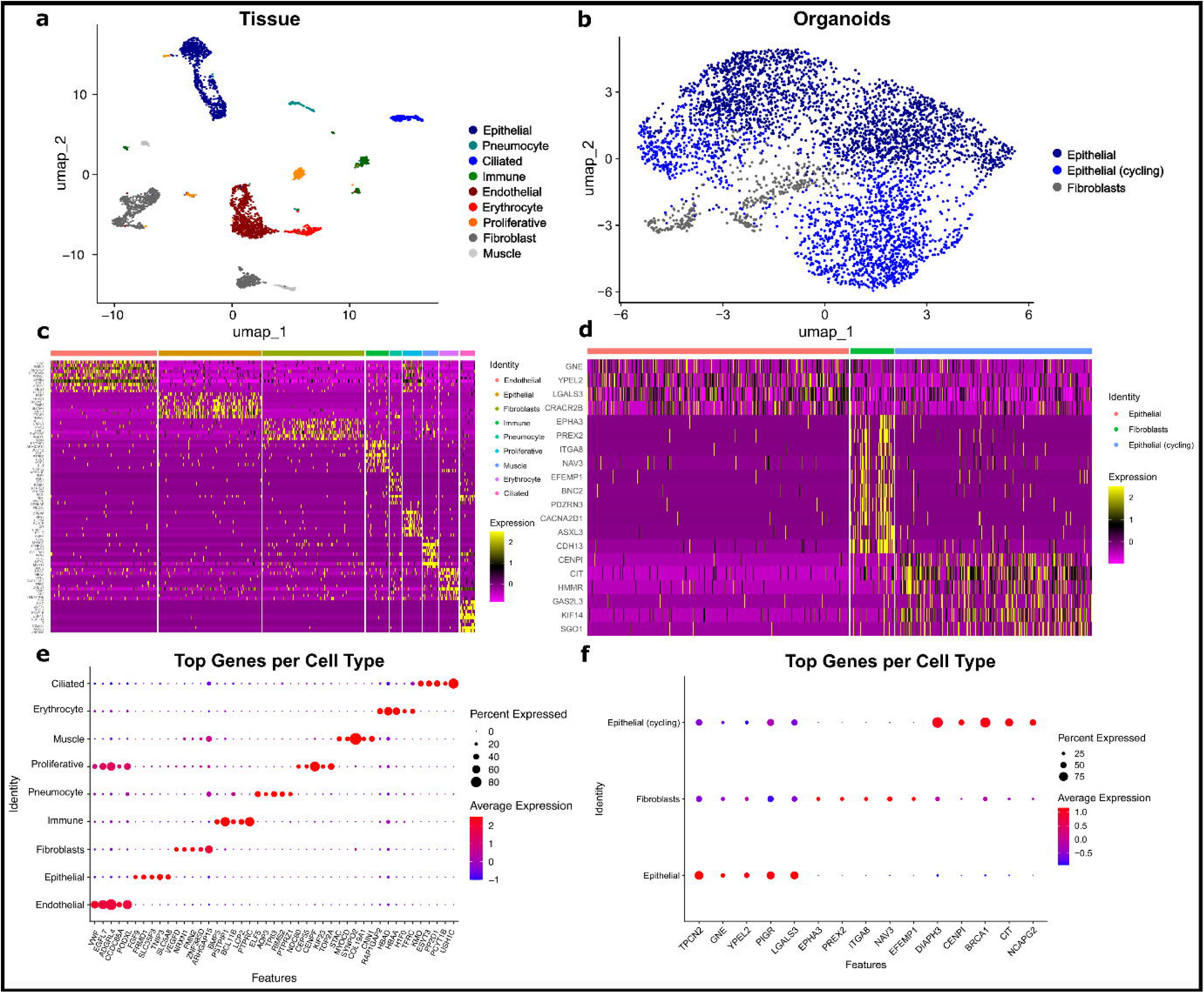
Identification of cell populations using snRNA-Seq analysis of chicken lung organoid and tissue of origin. UMAP projection of (**a**) chicken lung tissue and (**b**) organoids with clusters identified. Heatmaps of the top 10 genes expressed in each cluster in (**c**) tissue and (**d**) organoids. Dot plots indicate the top 5 genes associated with each cell population in (**e**) tissue and (**f**) organoids.

## Discussion

Adult stem cell-derived chicken lung organoids offer several advantages for *in vitro* modeling of viral pathogenesis and replication compared to current research strategies. Mammalian immortalized cells, such as Madin-Darby Canine Kidney (MDCK) cells, are commonly used to propagate and study avian influenza viruses (Tsai et al., 2019). However, studies have demonstrated significant differences in the replicative cycle of avian influenza viruses between avian and mammalian cells (G. Gabriel et al., 2007; Ganti et al., 2021). Furthermore, domestic chickens lack certain pattern recognition receptors, such as RIG-I, which play a key role in innate immune responses in other hosts of influenza viruses (Barber et al., 2010; Bertran et al., 2019). Galliformes utilize melanoma differentiation-associated gene 5 (MDA5) to promote type I interferon production and initiate an antiviral host response (Hornung et al., 2006; Karpala et al., 2011). These differences in viral sensing underscore the need for species-specific models. In the chicken lung organoids, the genetic precursor to MDA5 was present, while RIG-I was not.

Several chicken cell lines, including epithelial and mesenchymal lines, have been described (Esnault et al., 2011; Himly et al., 1998; Khatri et al., 2010; Zaffuto et al., 2008). While informative, these cell lines lack cellular heterogeneity, a key limitation for studying HPAIV strains known to infect various respiratory cell types (Rebel et al., 2011). Additionally, these lines are grown in two-dimensional monolayers, failing to replicate the three-dimensional architecture of lung tissue. In contrast, the chicken lung organoids described here provide a promising platform for studying chicken respiratory diseases, with a complex cellular composition resembling the avian lung epithelium.

ASC-derived organoids offer several advantages over iPSC-derived models, particularly in their rapid establishment. The ASC-derived lung organoids in this study showed growth within 24 hours post-isolation, while iPSC-derived models can require months to develop (Miller et al., 2019). Moreover, ASC-derived organoids recapitulate structures across both proximal and distal airways, whereas iPSC-derived models typically mimic specific lung regions (Han et al., 2022). These features, combined with their ability to reduce reliance on live animals, position organoids as an ethically and scientifically advantageous model (Carbone, 2011).

Despite lacking immune, stromal, and muscle cells found *in vivo*, histological comparisons demonstrated that the organoids retained key epithelial characteristics of the parent tissue. Transcriptomic analysis showed that genes of interest were similarly expressed between organoids and tissue samples, suggesting high fidelity in modeling chicken lung epithelial populations. While bulk RNA sequencing revealed differences between inflammatory and proliferative pathways, these results were expected because of the absence of immune components and presence of proliferative stem cells in the organoid samples, contrasting with intact immune systems in the tissues. Single-nuclei RNA sequencing identified epithelial cells and fibroblasts, further confirmed by pan-cytokeratin staining and downregulation of fibroblast gene markers (*COL1A1*, *COL1A2*) in organoids compared to tissue. Morphological and transcriptomic evidence suggests that the organoids presented here have a resemblance to the original respiratory epithelial compartments.

This study has limitations, including a small sample size (n = 3 donors) and the use of a single chicken breed. Future studies should explore organoids from other breeds and ages to capture the diversity of commercial chicken populations. Additionally, the effects of extended passages and thawing samples on genetic stability and cellular composition of the organoids should be further evaluated for functional assays. Further antibody optimization of immunofluorescence to confirm protein expression and spatial localization is also warranted. Additional analysis of single-nuclei RNA sequencing data is required to further differentiate cell types in the organoid samples, and single-cell RNA sequencing should be considered to identify cytoplasmic transcripts.

Organoids derived in this study have broad potential applications including infection with respiratory viruses to study viral pathogenesis, tropism, and replication dynamics, potentially predicting viral evolution. Of note, genes associated with inflammasome and NF-κB pathways were expressed in the organoids. The high mortality rate of HPAIVs in chickens underscores the need for such models to guide research on interventions. While preventative vaccines exist, they have not been allowed in the United States until recently. Additionally, as there is no treatment for avian influenza in poultry, recent evidence viral evolution demands further research into discovery of new preventions and treatments (Swayne & Kapczynski, 2016; Tseng et al., 2024). Human lung organoids have been used for similar purposes, and the strategies employed could be adapted to chicken lung organoids (Han et al., 2022; Salahudeen et al., 2020; S. Zhao et al., 2023).

With further regard for viral studies, organoids have been cultured from human lungs to model a variety of viruses, including avian and swine influenza strains, SARS-CoV-2, adenoviruses, and respiratory syncytial virus (Harford et al., 2022; Salahudeen et al., 2020; S. Zhao et al., 2023; J. Zhou et al., 2018). Previous studies have confirmed host tropism and viral replication in human organoids by determining viral loads, performing immunofluorescent staining, using single-cell RNA sequencing, and conducting transmission electron microscopy. This has helped to identify target cell populations and determine differences in infectivity between viral strains (Han et al., 2021; S. Zhao et al., 2023; J. Zhou et al., 2018). Furthermore, multiple studies have reported inhibition of viral entry and replication, indicating the potential use of lung organoids for screening antiviral treatments (Han et al., 2021; S. Zhao et al., 2023).

Hemagglutinin on avian influenza viruses binds to sialic acid receptors primarily found on the apical surface of respiratory epithelial cells, but the orientation of organoids is such that the apical side of the cell is facing inwards (C. Zhao & Pu, 2022; J. Zhou et al., 2018). Therefore, studies involving viral infection of human respiratory organoids have required the development of improved methods to access these receptors. The virus can be microinjected into the lumen of the organoids, or their polarization can be altered by either growing the cells as a monolayer on Transwell permeable supports or creating an apical-out orientation (Harford et al., 2022; Salahudeen et al., 2020; J. Zhou et al., 2018).

The successful establishment of chicken lung organoids suggests that similar methods could be applied to other avian species, such as waterfowl and raptors, in order to study the viruses in emerging and natural reservoir populations (Adlhoch et al., 2023; Nemeth et al., 2023). This is especially critical given the increasing number of spillover events and the potential for HPAIVs to evolve in ways that will enable human-to-human transmission (Burrough et al., 2024; de Vries & de Haan, 2023; Neumann & Kawaoka, 2024; Ulloa et al., 2023; WHO, 2025). Therefore, developing a better understanding of the disease in birds may help us better understand the viral changes leading to mammalian, including human, adaptation.

Beyond influenza, the model may be useful for investigating other poultry respiratory pathogenic infections that illicit similar antimicrobial immune responses, including Newcastle disease, infectious bronchitis, chlamydiosis, and colibacillosis (Boodhoo et al., 2016; Glisson, 1998; Kameka et al., 2014; Yehia et al., 2023). Organoids have already been used to model bacterial infections in other systems, and similar approaches could be applied here (Iakobachvili et al., 2022; Tang et al., 2022).

Future directions include infecting the organoids with various influenza viruses to determine how the extent of the infection dynamics is comparable to other existing *in vitro* and *in vivo* models. Another point of interest would be the development of organoid-immune cell co-cultures to better model host-pathogen interactions. These systems, though currently focused on cancer research, could provide significant insights into immune responses during respiratory infections (Bar-Ephraim et al., 2020; Cattaneo et al., 2020; Chakrabarti et al., 2021). Co-culture models could also advance our understanding of diseases such as Marek’s disease (*Mardivirus gallidalpha2*, *Gallid alphaherpesvirus 2*), which targets T lymphocytes following epithelial infection and can, in some cases, lead to lymphoma (Boodhoo et al., 2016). In conclusion, the development of ASC-derived chicken lung organoids represents a significant advancement in avian respiratory research. These models have potential applications in developmental biology, infectious diseases, and the study of avian influenza, addressing critical needs in both veterinary and public health fields.

## Data availability

The stranded mRNA raw RNA-seq reads and snRNA-seq reads generated and analyzed in this study are available in the Gene Expression Omnibus (GEO) under accession codes GSE291341 and GSE291342.

## Code availability

Bioinformatic scripts are available on GitHub (https://github.com/h-nicholson/Chicken_Lung_Organoids) and (https://doi.org/10.5281/zenodo.15009183).

## Acknowledgments

We thank Mohamed Elbadawy and the SMART Lab members for their help with the samples. Additionally, we thank William Bastian for his assistance in the snRNA-seq analysis. We appreciate the timely processing of samples at the University of Georgia Histology Laboratory. We wish to thank Mary Ard of Georgia Electron Microscopy for processing and imaging TEM samples.

## Competing interests

K. Allenspach is a co-founder of LifEngine Animal Health and 3D Health Solutions. She serves as a consultant for Ceva Animal Health, Bioiberica, LifeDiagnostics, Antech Diagnostics, Deerland Probiotics, Christian Hansen Probiotics, Purina, and Mars. J.P. Mochel is a co-founder of LifEngine Animal Health (LEAH) and 3D Health Solutions. Dr. Mochel is a consultant for Ceva Animal Health, Ethos Animal Health, LifEngine Animal Health and Boehringer Ingelheim. C. Zdyrski is the Director of Research and Product Development at 3D Health Solutions. Other authors do not have any competing interests to declare.

## Funding

This project was partially funded by startup funds provided by the University of Georgia, the University of Georgia College of Veterinary Medicine, and by the U.S. National Science Foundation project ID FP00029141, Award ID 2344946.

## Supplemental Materials

**Supplemental Table 1**: Composition of Shipping Media, including reference numbers and component concentrations.

**Supplemental Table 2**: Media composition of CMGF+ R/G (Complete Media with Growth Factors + ROCK inhibitor and CHIR99021 (GSK-3β inhibitor)), with distributor reference numbers and concentrations.

**Supplemental Table 3**: Information about each donor animal, including breed and age of the chickens. Additionally, information is provided about each organoid culture, which includes the number of days in culture, number of passages, preservation methods, and contamination testing results.

---

**Supp. Fig. 1:**
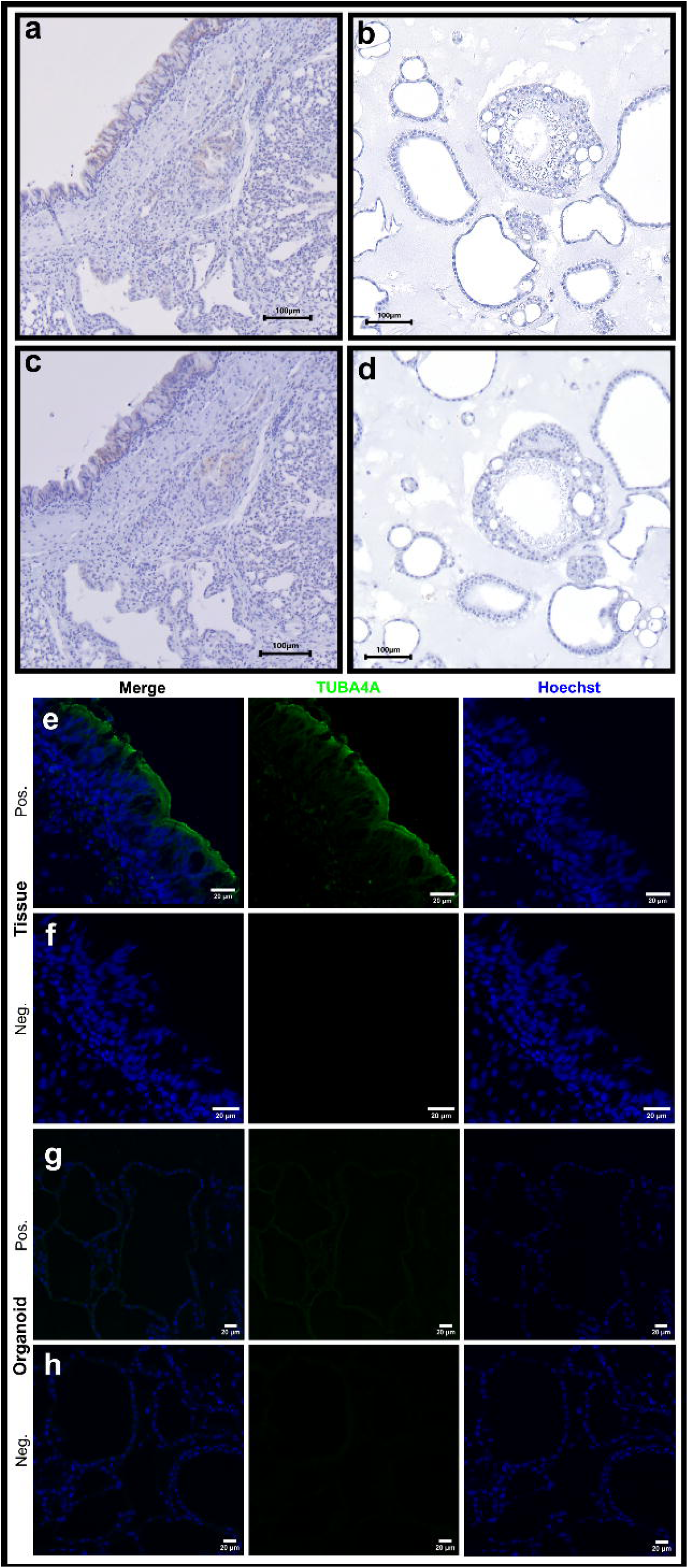
Negative controls for immunohistochemistry and immunofluorescence staining. Negative control of immunohistochemical pan-cytokeratin staining of (**a**) chicken lung tissues and (**b**) organoids for pan-cytokeratin. Negative control of immunohistochemical TTF-1 staining (**c**) chicken lung tissues and (**d**) organoids. Immunofluorescent staining of chicken lung tissues with an antibody against (**e**) acetylated alpha-tubulin (TUBA4A), as well as staining nuclei (Hoechst) and (**f**) a negative control. Immunofluorescent staining of chicken lung organoids with (**g**) TUBA4A and (**h**) a negative control. Scale bars are in μm.

## Notes

https://github.com/h-nicholson/Chicken_Lung_Organoids

https://doi.org/10.5281/zenodo.15009183

